# Difference in determinants and changing patterns of plant community composition with and without grazing across semi-arid and arid regions in Mongolian steppe

**DOI:** 10.1101/2022.12.24.521851

**Authors:** Naohiro I. Ishii, Issei Nishimura, Yulan Qi, Gantsetseg Batdelger, Maiko Kagami, Gaku Takimoto, Takehiro Sasaki

## Abstract

Aridity, edaphic variables related to aridity, and livestock grazing are major drivers of plant community composition across dryland grasslands. With accounting these factors, little is known about differences in determinants and changing patterns along aridity gradients with different average of aridity. Thus, the comparative investigations of communities without and with grazing in semi-arid and arid regions are suitable to clarify the difference in compositional responses along aridity gradients between potential vegetations and degraded ones with grazing. We investigated compositional changes of communities without/with grazing in semi-arid (north) /arid (south) regions across Mongolia. The compositional changes based on Bray-Curtis dissimilarity were investigated by generalized dissimilarity modeling, including geographic distance, aridity, soil pH, and sand and clay contents as independent variables. The determinants and changing patterns of community composition were compared among the four groups. Aridity had significant impacts on community composition regardless of the regions and the absence/presence of grazing. However, without the dependency on grazing, the difference in response patterns was observed between the regions. The compositional change was steeper especially at the upper edge of aridity rather than the lower edge in the arid region. This indicates the vulnerability of plant communities to aridity shifts due to future climate change in desert steppe of Mongolia. In addition, regardless of the regions, the effects of soil pH on community composition were eliminated by grazing. Because soil pH indirectly affected by aridity shifts can have impacts on community composition of potential vegetations without grazing, the long-term monitoring of vegetation dynamics needs observations of both of communities without and with grazing.

## Highlights

- Determinants and changing patterns in plant community composition of Mongolian steppe along aridity gradients in semi-arid and arid regions were investigated with the consideration of the effects of aridity and edaphic variables for communities with the absence/presence of livestock grazing, respectively.
- Although aridity had significant impacts on community composition regardless of the regions and the absence/presence of grazing, the difference in response patterns was observed between the regions, which did not depend on grazing.
- Regardless of the aridity levels, the effects of soil pH on community composition were eliminated by grazing.

## Introduction

Plant community composition is a central factor determining the impacts of climate change and anthropogenic disturbances on ecosystem structures and functioning (Chapin et al., 2000). Across dryland ecosystems, aridity and livestock grazing are the most influential factors on these facets via the changes in plant community composition (Maestre et al., 2016).

Berdugo et al. (2020) reported abrupt shifts in the ecosystem structures (e.g. plant cover and species richness) along a broad gradient of aridity. Additionally, aridity affects not only water availability but also edaphic variables through long-term feedback from vegetations (Berdugo et al., 2020; Delgado-Baquerizo et al., 2013; Moreno-Jiménez et al., 2019) Because edaphic variables can have impacts on the patterns of community composition across dryland grasslands (Sasaki et al., 2008b; Zemmrich et al., 2010), the consideration of both of aridity and edaphic variables associated with aridity is required to clarify community assembly in drylands. However, little is known about the consistency in patterns of changes in community composition among different aridity gradients with this consideration. Understanding consistency or inconsistency in compositional changes across different gradients of aridity can provide implications for the impacts of aridity shifts under climate change on plant communities and their functioning in dryland ecosystems.

Grassland vegetations are widely used for livestock grazing to support secondary production in drylands. Numerous studies have revealed the impact of grazing on community composition mediated by selective herbivory (Díaz et al., 2007; Li et al., 2022; Sasaki et al., 2008a). The grazing impacts on community composition can differ according to differences in water availability. Previous studies have demonstrated that the relative magnitude of grazing impacts on vegetation is relatively low in more arid regions (Herrero-Jáuregui and Oesterheld, 2018; Price et al., 2022). However, regardless of aridity levels, community composition can be drastically altered and homogenized by high intensity of grazing, namely overgrazing (Ahlborn et al., 2020; Sasaki et al., 2009b, 2008a; Wang et al., 2017). Thus, the patterns of changes in community composition along aridity gradients in grazed grasslands can be confounded depending on grazing. To infer how the changes of community composition along different aridity gradients will be driven by climate change, the comparisons of compositional changes are essential with the consideration of the interaction between aridity levels and grazing. Notably, these comparisons have not been conducted.

Here, we investigated the patterns of changes in plant community composition along aridity gradients with different levels of average aridity (semi-arid and arid regions) in the Mongolian steppe depending on the absence/presence of grazing (without/with the long-term exclosure of livestock). Mongolia covers a vast area of the eastern part of the Eurasian Steppe. In Mongolia, the climate is spatially characterized by broad variations in aridity associated with temperature and precipitation (Vandandorj et al., 2017). Suzuki et al. (2022) demonstrated that aridity was the important driver of plant community composition across the whole range of Mongolia. Thus, we tested the research questions below mentioned to provide implications into the vulnerability of the Mongolian steppe communities and the long-term monitoring of vegetation dynamics to climate change: 1) whether and how the effects of aridity and edaphic variables on community composition differ between semi-arid and arid grasslands, and 2) does grazing alter these effects.

## Materials and methods

Across Central Mongolia, ranging from semi-arid to arid regions, over 600 km, 12 study sites were selected at the meteorological sites of the Mongolian Information and Research Institute of Meteorology, Hydrology, and Environment (IRIMHE) (Table 1, and Figure S1). We aimed to ensure that the study area covered a range of vegetation types within the rangeland ecosystems of Mongolia. Central Mongolia has the typical climate of Central Asia, which is characterized by hot and short summers with approximately 70 % annual precipitation and cold winters, including extreme climate events such as drought. Based on the climate data between 1978 and 2017 provided by IRIMHE, the mean annual precipitation ranged from 97 mm at Tsogt-Ovoo to 333 mm at Darkhan; the mean annual temperature ranged from −0.8 ℃ at Ugtaal to 7.6 ℃ at Khanbogd. Thus, there is a wide gradient of aridity from north to south regions (Figure S1), which supports three main vegetation types: forest steppe, steppe, and desert steppe (Table 1).

**Table 1.**
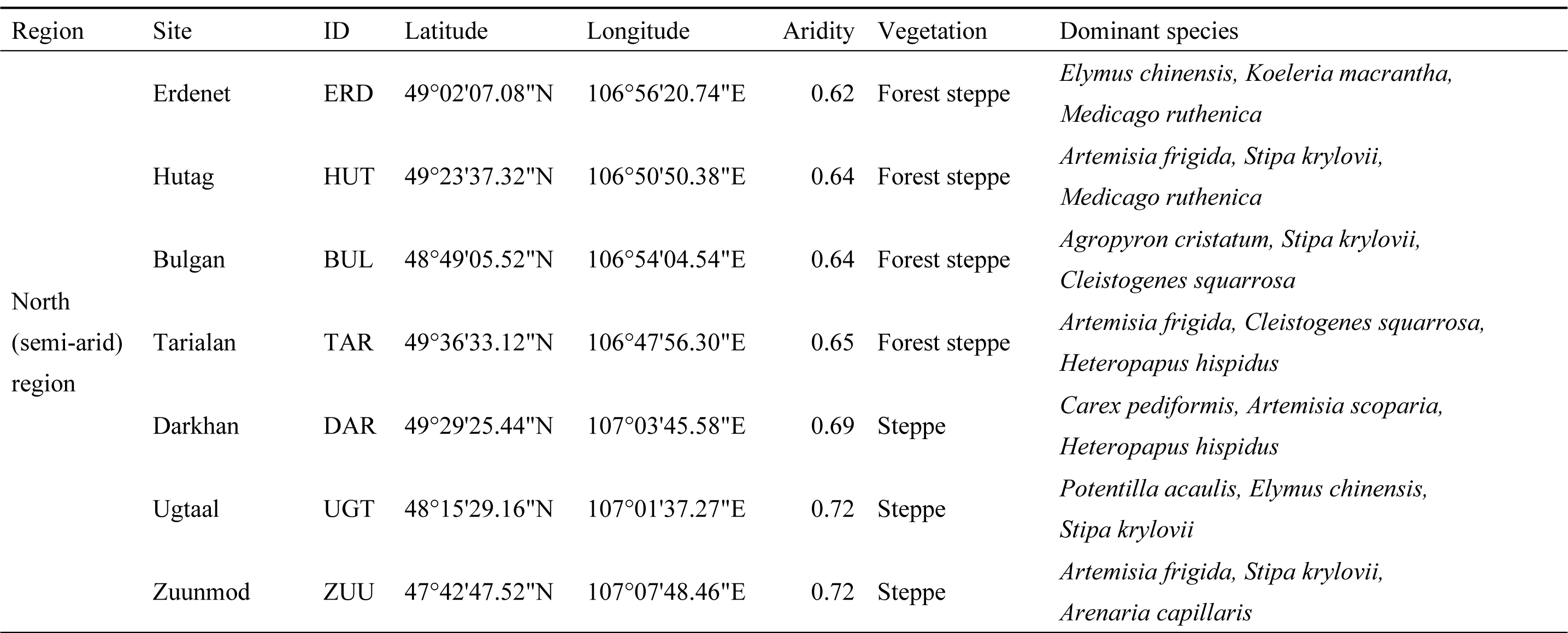
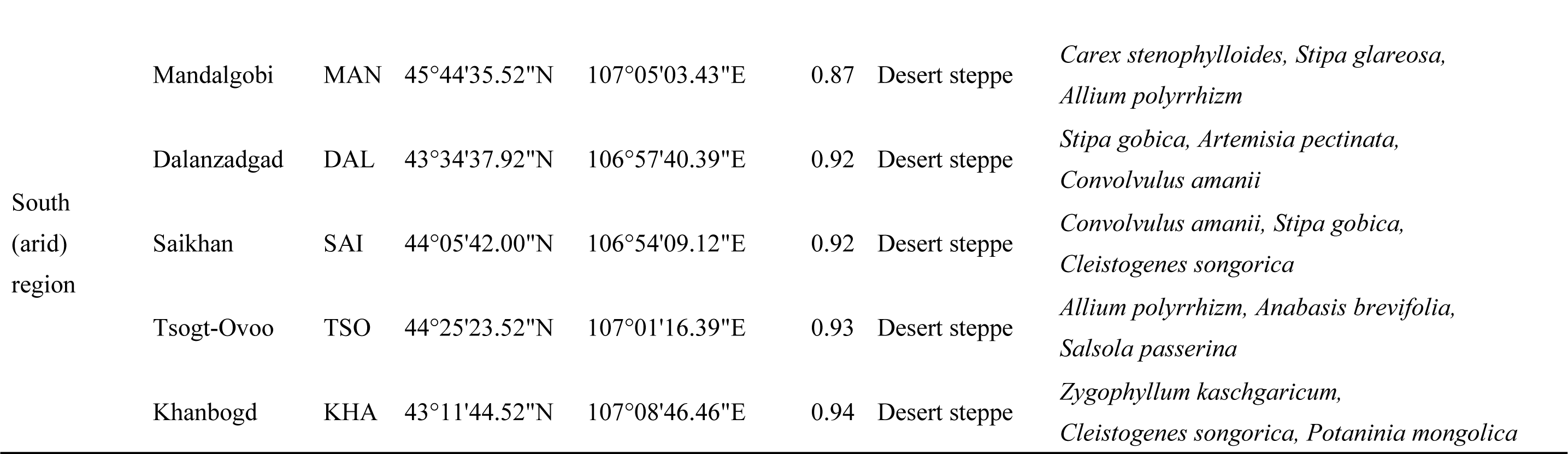
Locations, aridity, vegetation type, and dominant species of the 12 study sites. Aridity is calculated as 1 – (precipitation/potential evapotranspiration).

We set three 5 × 5 m blocks, which included three 1 × 1 m plots, at inside and outside of meteorological sites (50 × 50 m), to observe the heterogeneity at both block- and site-level scales. All the sites had ungrazed and grazed replicates. Ungrazed ones were exclosed for at least 50 years to avoid livestock grazing. Because the distances between ungrazed and grazed replicates in each site varies among sites, from a minimum of 25 m to a maximum of 3 km, the comparison of community composition between ungrazed and grazed replicates in each site was not performed in this study. In each plot, vegetation was sampled by identifying all species rooting in each plot, and absolute cover (%) of each species was recorded from July to August 2022. Then, the vegetation data were pooled for each block. Simpson diversity index, which reflects the diversity index mainly based on relative abundance of dominant species, was almost saturated when the vegetation data of more than two plots was used for the calculation of Simpson index in each site (Figure S2).

Aridity, which is calculated as [1–precipitation/potential evapotranspiration], based on climatic data for the 1970–2000 period was obtained from the Global Aridity Index and Potential Evapotranspiration Climate Database v3 (Trabucco and Zomer, 2022). The edaphic variables were investigated based on the measurement of a soil sample in each block: total nitrogen content in cg/kg; soil organic carbon content in dg/kg; soil pH in H_2_O; clay, sand, and silt content mass fraction in %. Soil cores with 5 cm in diameter and 10 cm in length in three plots were pooled by each block, and used for the measurement of edaphic variables. Because of the high level of correlations (absolute value of a correlation > 0.6) between aridity, and total nitrogen and soil organic carbon, and between sand and silt content, we used the aridity, soil pH, sand content, and clay content as the environmental variables in the subsequent analysis.

Nonmetric multi-dimensional scaling (NMDS) based on Bray-Curtis dissimilarity was performed for the whole dataset of vegetation using the function ‘metaMDS’ of the R package ‘vegan.’ The dimensional reduction to two dimensions was performed because the stress value (0.093) was lower than 0.2 (Kruskal, 1964). In addition, the clustering based on Ward’s algorithm was conducted using the Bray-Curtis dissimilarity matrix.

We performed the analysis independently for the following four groups: the groups from the north (semi-arid) and south (arid) regions in Mongolia without/with grazing (north-ungrazed, north-grazed, south-ungrazed, and south-grazed). To assess the difference in community composition among the study sites, Bray-Curtis dissimilarity was calculated for each group. The abundance-based dissimilarity was then partitioned into balanced variation in abundance and abundance gradient using the function ‘beta.multi.abund’ of the R package ‘betapart’ (Baselga, 2017). This partitioning informs relative contributions of species turnover (balanced variation) and nestedness (abundance gradient) to community composition difference. Moveorver, to identify factors influencing community compositions, we applied generalized dissimilarity modeling (GDM; Ferrier et al. (2007)). GDM is a matrix regression technique that permits non-linear responses of compositional dissimilarity to spatial or environmental gradients. This approach has an additional advantage to depict the rate of compositional shift at any point along the gradient of a variable because GDM fits monotonically increasing flexible *I*-spline functions for each independent variable. When a *I*-spline was plotted, the maximum height depicts the magnitude in an effect of a given variable on compositional dissimilarity (partial dissimilarity) with holding all the other independent variables constant. The first derivatives (slopes) of a spline indicates the rate of compositional shift at any point along the gradient of a variable. Using the function ‘gdm’ of the R package ‘gdm’, we fitted full GDMs with pairwise geographical distance and the selected environmental variables as independent variables for each group. These models were fitted with the three *I*-spline basis functions per variable, and regression coefficient at each point (lower edge, mid-point, and upper edge) along an independent variable was estimated. The uncertainty for each GDM plot (standard deviation) was calculated using the function ‘plotUncertainty’ with 1,000 bootstrap iterations. Additionally, we fitted GDMs with a single independent variable. From these models, the significance of each variable was calculated using Monte Carlo permutation by the function ‘gdm.varImp.’ We investigated 1) whether and how the effects of aridity and edaphic variables on community composition differ between semi-arid and arid grasslands, based on the comparisons in the results of GDMs between the north and south regions. In addition, the comparisons in the results of GDMs between the ungrazed and grazed communities assessed that 2) does grazing alter these effects. To check correlations between the matrices of distance (geographical distance and differences in the selected environmental variables), the calculation of Mantel’s correlation coefficient based on Spearman correlation coefficient and Mantel test for each pair of the variables were performed.

## Results

The result of NMDS and clustering showed that there were mainly two groups of community composition along the first axis (Figure S3–S5). These two groups corresponded to the communities in the north (forest steppe) and south (steppe and desert steppe) regions in Central Mongolia (Figure S5). The partitioning of Bray-Curtis dissimilarity exhibited the large contribution with more than 90% of balanced variation in abundance (species turnover) rather than abundance gradient (nestedness) (Table S1). GDMs revealed that the significant effects of geographic distance and aridity on the variation of community composition across all the groups (Table 2). On the other hand, only in the ungrazed communities of the both regions, the impacts of soil pH on community composition were significant. We observed the similar response patterns of community composition (partial Bray-Curtis dissimilarity) to geographic distance and soil pH in the both regions (Table 2, and Figure 1). However, the different response patterns to aridity were visually observed between the north and south regions: the steeper slope in the lower edge of aridity and saturated response in the upper edge in the north region, and the steeper response in the mid-point and upper edge in the south region. This pattern was not depended on the absence/presence of grazing. Mantel correlation coefficients between geographic distance and aridity difference in the north and south regions were similarly high, 0.748 and 0.790, respectively (Table S2).

**Figure 1.**
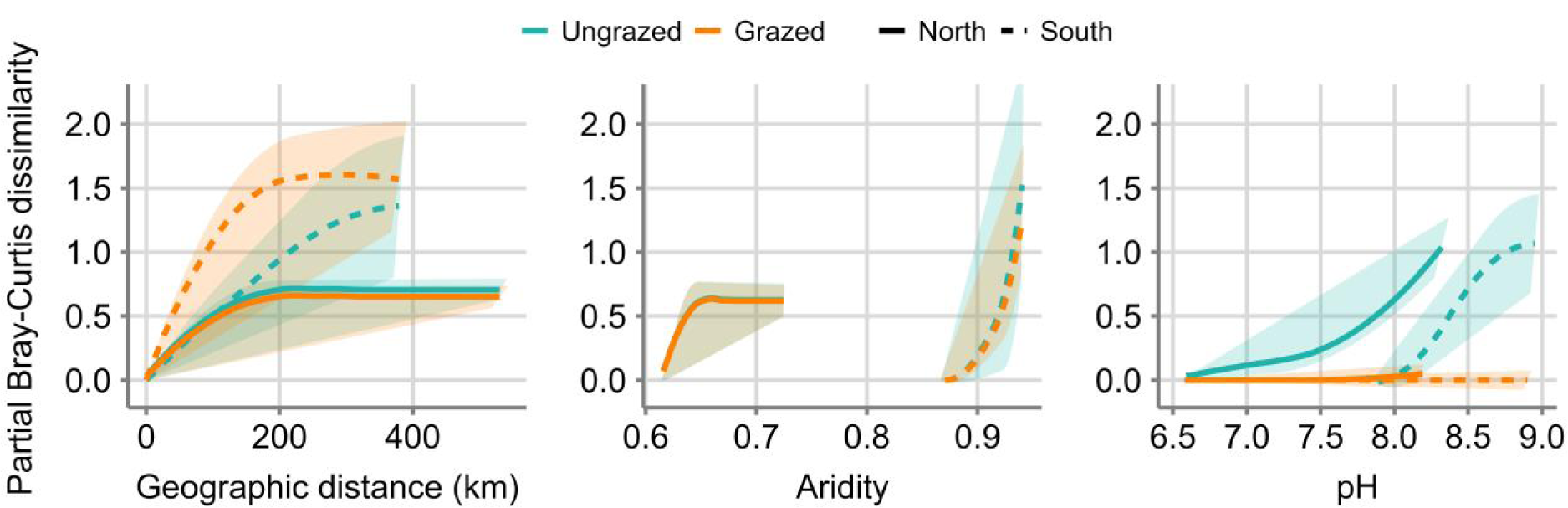
Relationships between partial Bray-Curtis dissimilarity and geographic distance, aridity, and soil pH based on generalized dissimilarity modeling (GDM). The estimated splines and their confidence intervals are denoted.

**Table 2.**
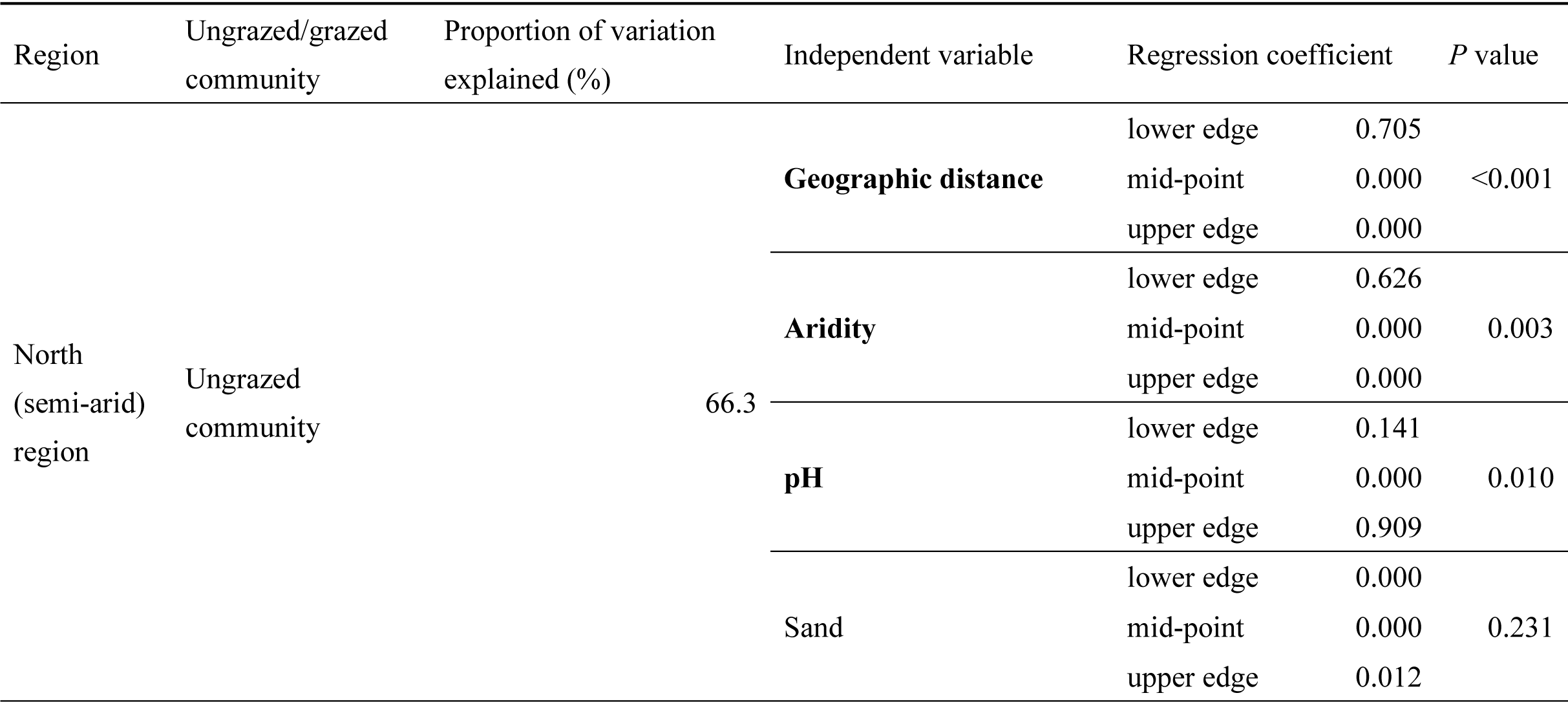
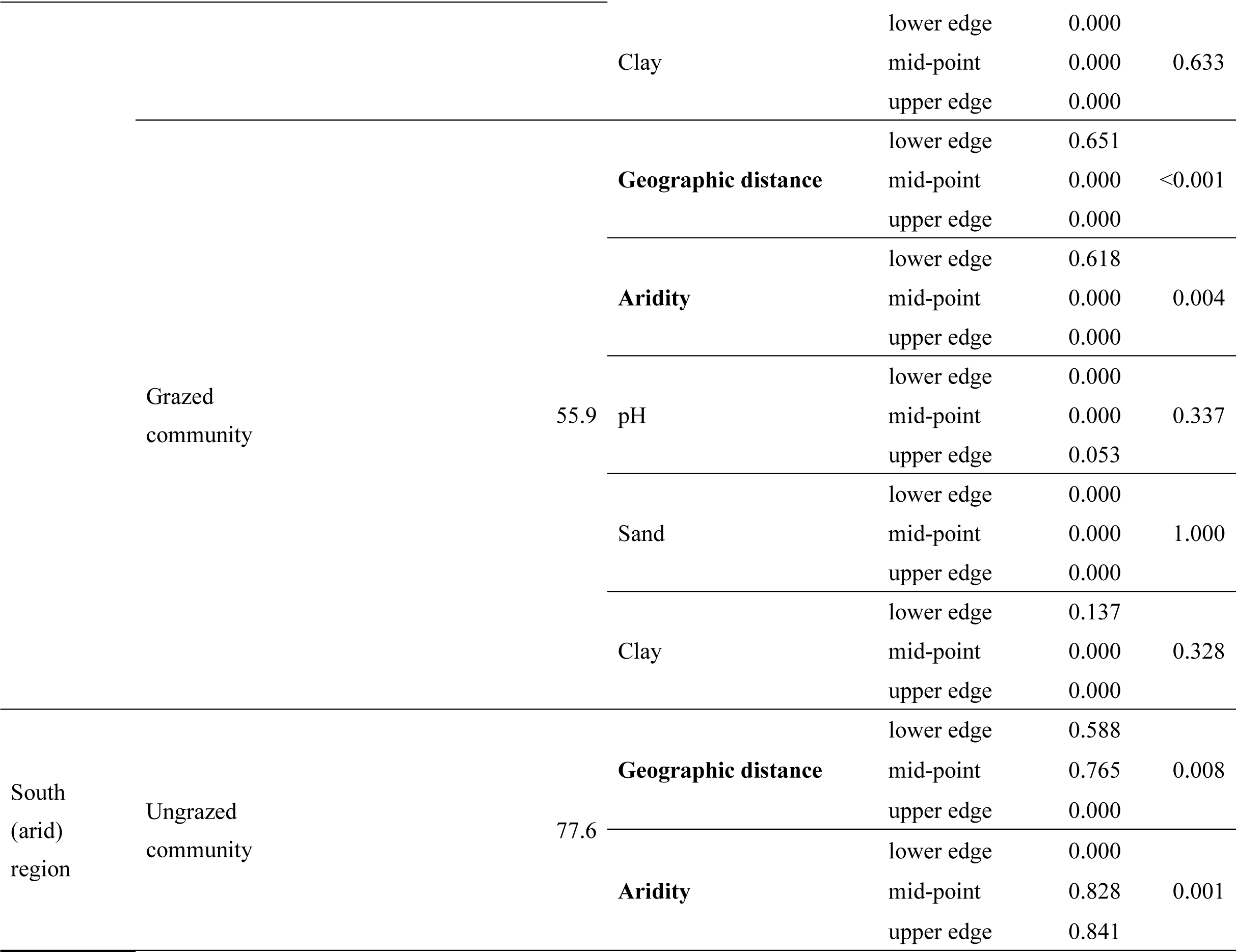
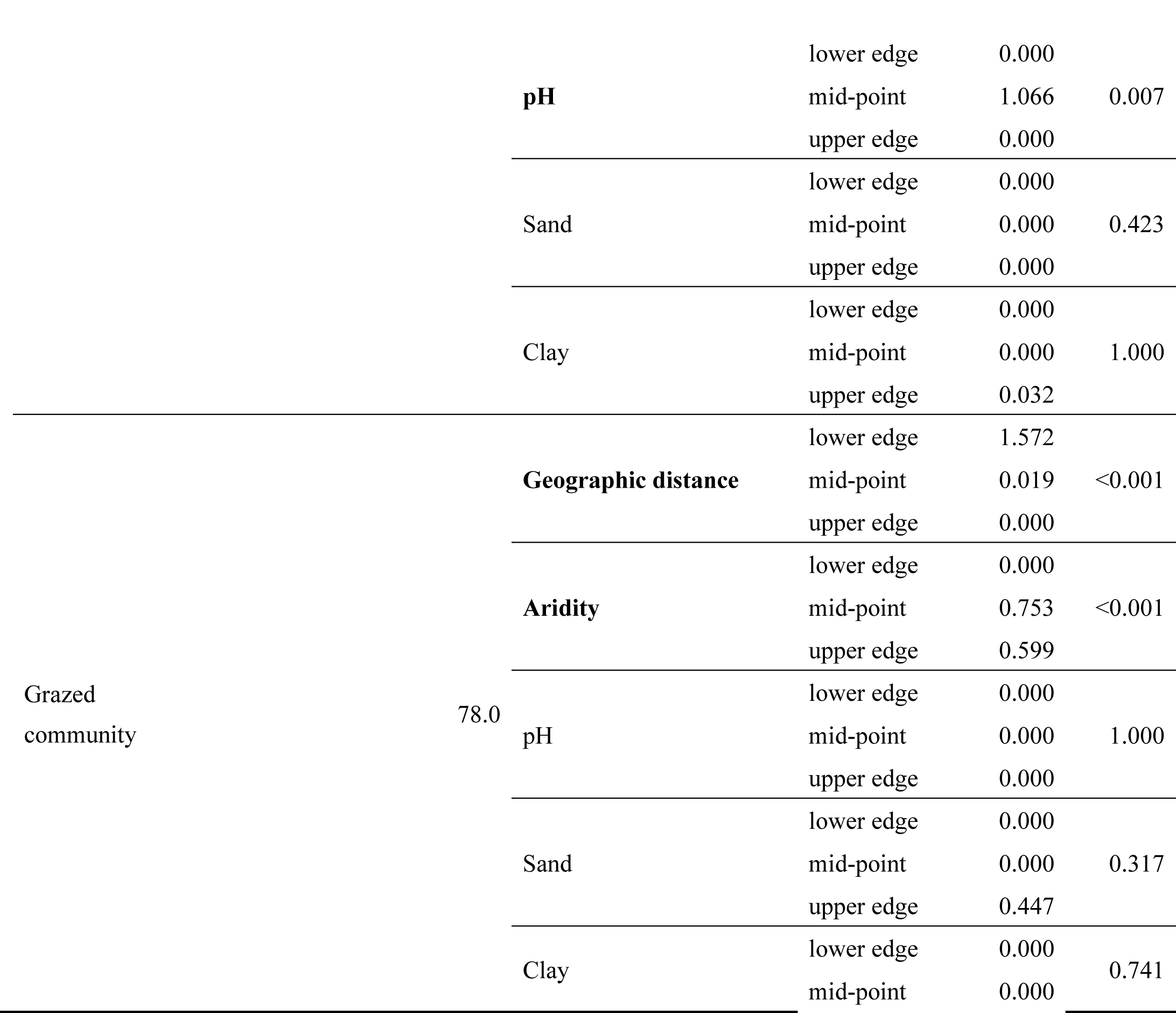
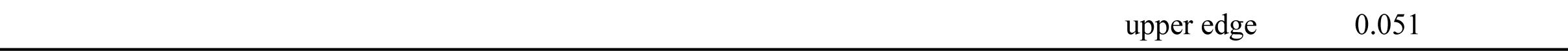
Regression coefficients and their significance of geographic distance, aridity, and soil pH based on generalized dissimilarity modeling (GDM) in ungrazed and grazed communities of the north and south regions. Regression coefficients on a spline (in Figure 1) at the lower edge, mid-point, and upper edge of each independent variable in each group (north-ungrazed, north-grazed, south-ungrazed, and south-grazed) are shown. Bold characters denote statistically significant variables.

## Discussions

Although aridity had significant impacts on community compositions in both of the semi-arid and arid regions of Mongolia (Table 2), we firstly demonstrated the difference in response patterns along the aridity gradients between these regions regardless of grazing (Figure 1). Thus, grazing did not confound the aridity impacts on community composition. The changes in community composition in the south and arid region was abrupt especially at the upper edge of aridity (Figure 1). This pattern might be mainly attributed to the relatively high level of environmental filtering and species turnover because of abiotic and biotic factors by high level of aridity. Because of the physiological limitations associated with aridity, the dominance of stress-tolerant species and exclusion of non-tolerant species can often be observed in dryland ecosystems (Berdugo et al., 2019; Grime, 1977; Le Bagousse-Pinguet et al., 2017). Previous studies have also demonstrated that high aridity contributes to the severe limitations of not only water availability but also soil nutrient availability, such as nitrogen, carbon, and micronutrients, via long-term feedback from vegetations (Berdugo et al., 2020; Delgado-Baquerizo et al., 2013; Moreno-Jiménez et al., 2019), resulting in severe environmental filtering by the abiotic factors associated with aridity. As a biotic factor, species turnover via competitive exclusion by highly adapted species in severely arid environments (Berdugo et al., 2019; Grime, 1977; Macarthur and Levins, 1967; Michalet et al., 2014) might attribute to the observed response patterns of community composition in this study. Increasing aridity can lead to the dominance of aridity specialists and an increase in the relative importance of competitive exclusion by the superior specialists, rather than facilitation (Berdugo et al., 2019). Some of previous studies have indicated that negative effects on water availability by aridity specialists (e.g. shrub species) can cause competitive exclusion in dryland ecosystems (Butterfield et al., 2016; O’Brien et al., 2017). In this study, one or two shrub species were dominant or predominant in the sites with the highest aridity (e.g., *Zygophyllum kaschgaricum*, *Potaninia mongolica*, *Anabasis brevifolia*, and *Salsola passerina*; Table 1) In addition, the relative abundance of these shrub species was positively correlated with aridity (Table S3), which might support the competitive exclusions by shrub species.

Our study also emphasized that the elimination of the effects of soil pH on grassland community composition across the semi-arid and arid regions by grazing (Table 2, and Figure 1). Although the relationships between species richness and soil pH, which is related to nutrient availability, were negative or absent in semi-arid and arid regions (Palpurina et al., 2017), the community composition including relative abundance of component species only without grazing was significantly affected by soil pH in both of the semi-arid and arid regions. This indicated that the grazed communities lost some variations of community composition associated with spatial heterogeneity of soil pH. In a coastal dune community, grazing removed the impact of soil pH on community composition (Tahmasebi Kohyani et al., 2008). In addition, grazing drastically decreased the number of species of which abundance were strongly affected by soil pH in a limestone plant community (Amezaga et al., 2004). Selective herbivory by livestock can lead to communities with biased life history and functional traits (e.g. annual and small-height plants) (Díaz et al., 2007), resulting in community composition less sensitive to soil pH via the reduction in diversity of responses to nutrient heterogeneity among species.

Our study indicated the vulnerability of plant communities to aridity shifts due to future climate change, especially in the upper edge of aridity range of Mongolia, namely desert steppes. Because accurate threshold detections for community composition along the aridity gradients (Berdugo et al., 2020) could not be performed in this study, future studies should use continuous and intensive vegetation sampling across the whole range of the Eurasian steppe and other regions. As a limitation in this study, there is no observation of temporal variability of vegetation and aridity. For example, Sasaki et al. (2009a) observed the inter-annual variations of plant community composition in the Mongolian steppe. Then, we propose the temporal observations of the relationships between community composition, aridity, and edaphic variables along multiple aridity gradients as a next step for understanding community assembly related to aridity, and its spatial and temporal variability. As another important aspect of this study, we raised the elimination of the impacts of soil pH, which is positively correlated with aridity globally (Moreno-Jiménez et al., 2019), on community composition by grazing. Because soil pH can have impacts on community composition of potential vegetations via indirect effects of aridity shifts, our study empathized the importance of not only grazed communities but also ungrazed ones for the long-term monitoring regarding vegetation dynamics exposed to climate change.

## Ethics approval

Not applicable.

## Consent to participate

All the authors approved participation in this study.

## Consent for publication

All the authors approved the publication of this study.

## Data availability

Data are available from the corresponding author upon reasonable request.

## Code availability

Codes are available from the corresponding author upon reasonable request.

## Author Contributions

N.I.I., G.B., G.T., M.K., and T.S. designed the study. All the authors contributed to the field surveys. N.I.I. performed the analyses and wrote the first draft of the manuscript. All the authors have contributed to the revisions.

## Acknowledgments

We would like to thank our laboratory colleagues for their help with the analyses.

## Supplementary information

**Figure S1.**
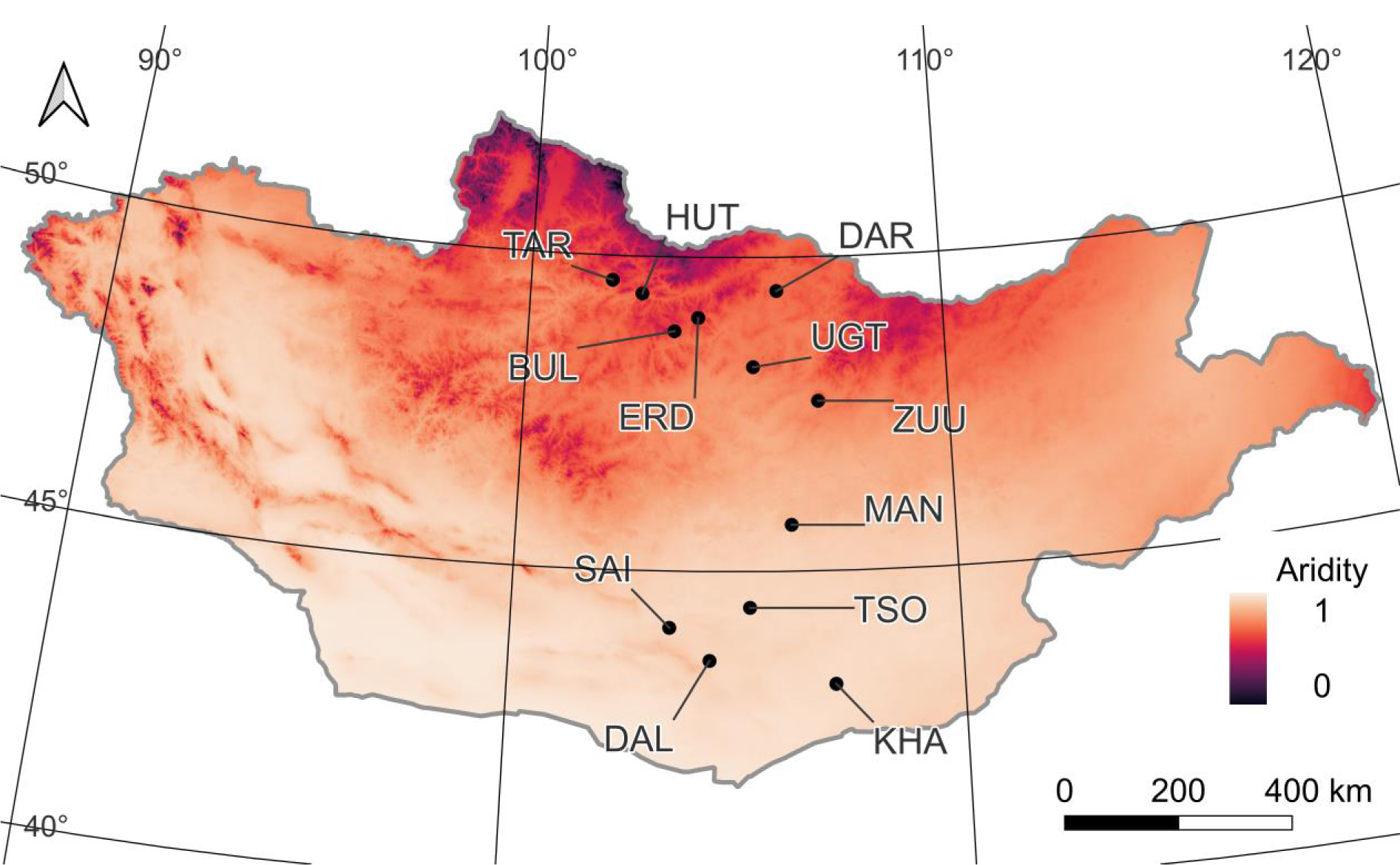
Spatial distribution of the 12 study sites and aridity in Mongolia. Aridity is calculated as 1 – (precipitation/potential evapotranspiration).

**Figure S2.**
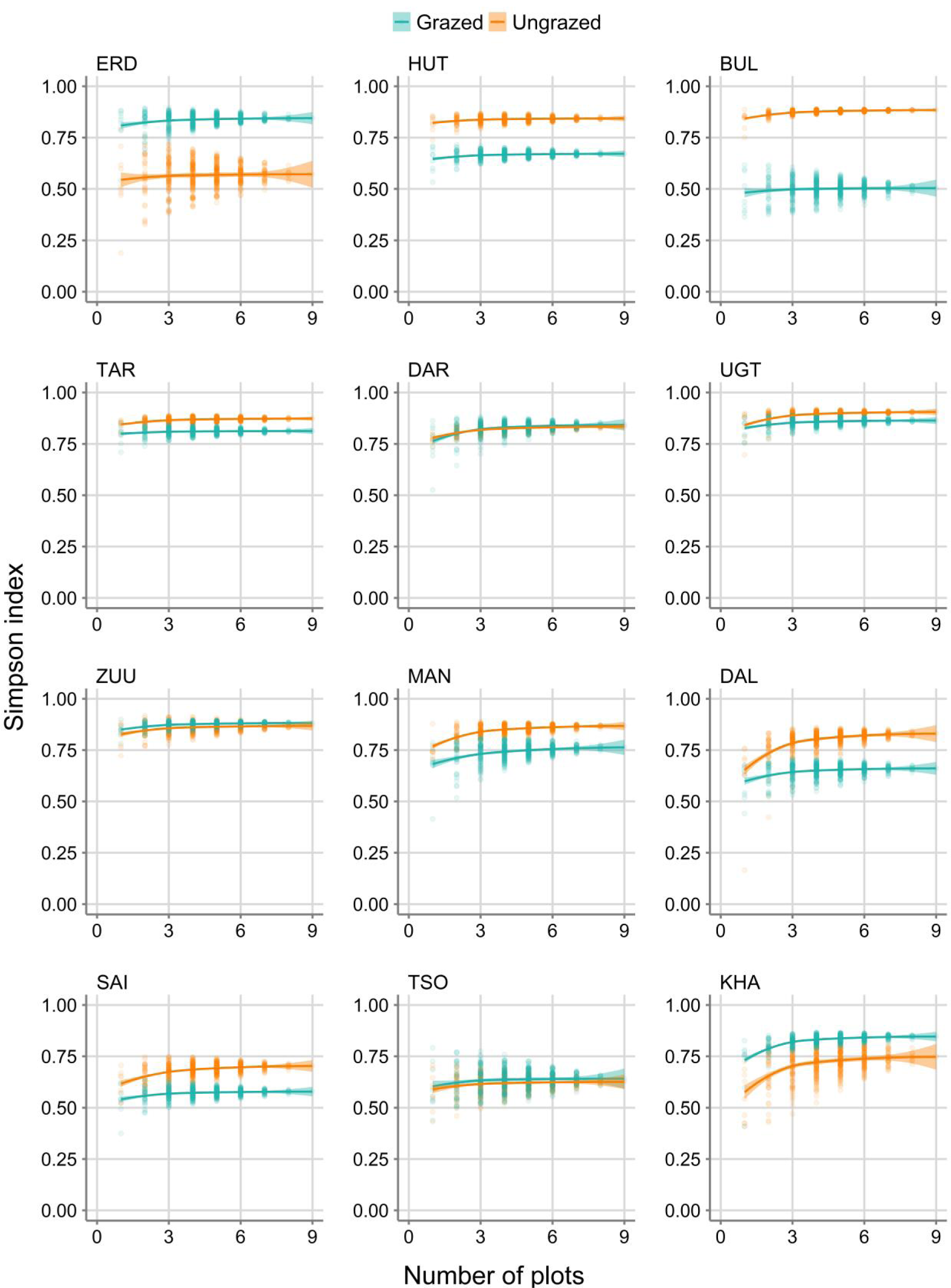
Relationships between number of plots and Simpson diversity index of ungrazed and grazed communities in each site.

**Figure S3.**
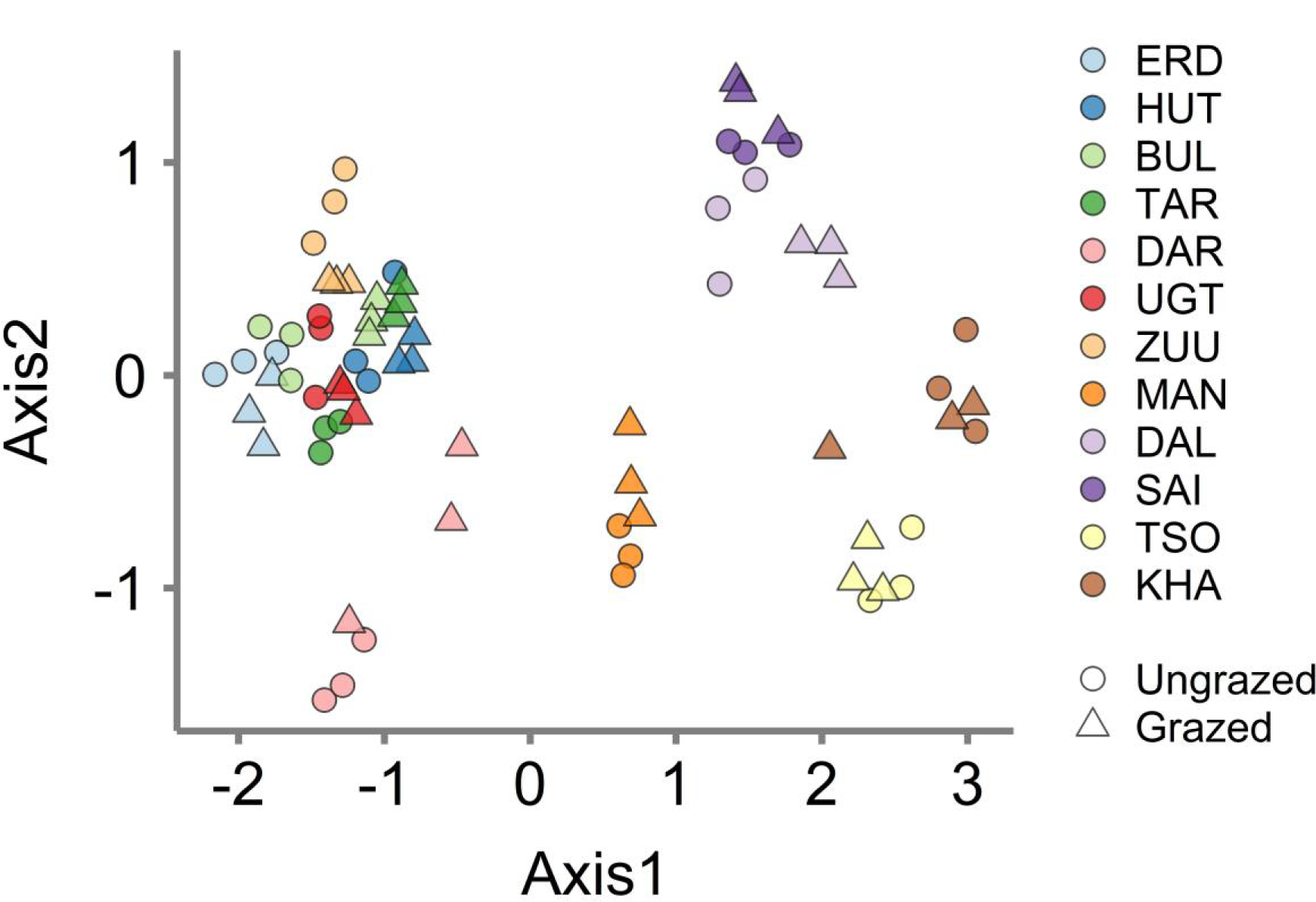
Result of nonmetric multi-dimensional scaling (NMDS) based on Bray-Curtis dissimilarity of ungrazed/grazed communities in the north and south regions.

**Figure S4.**
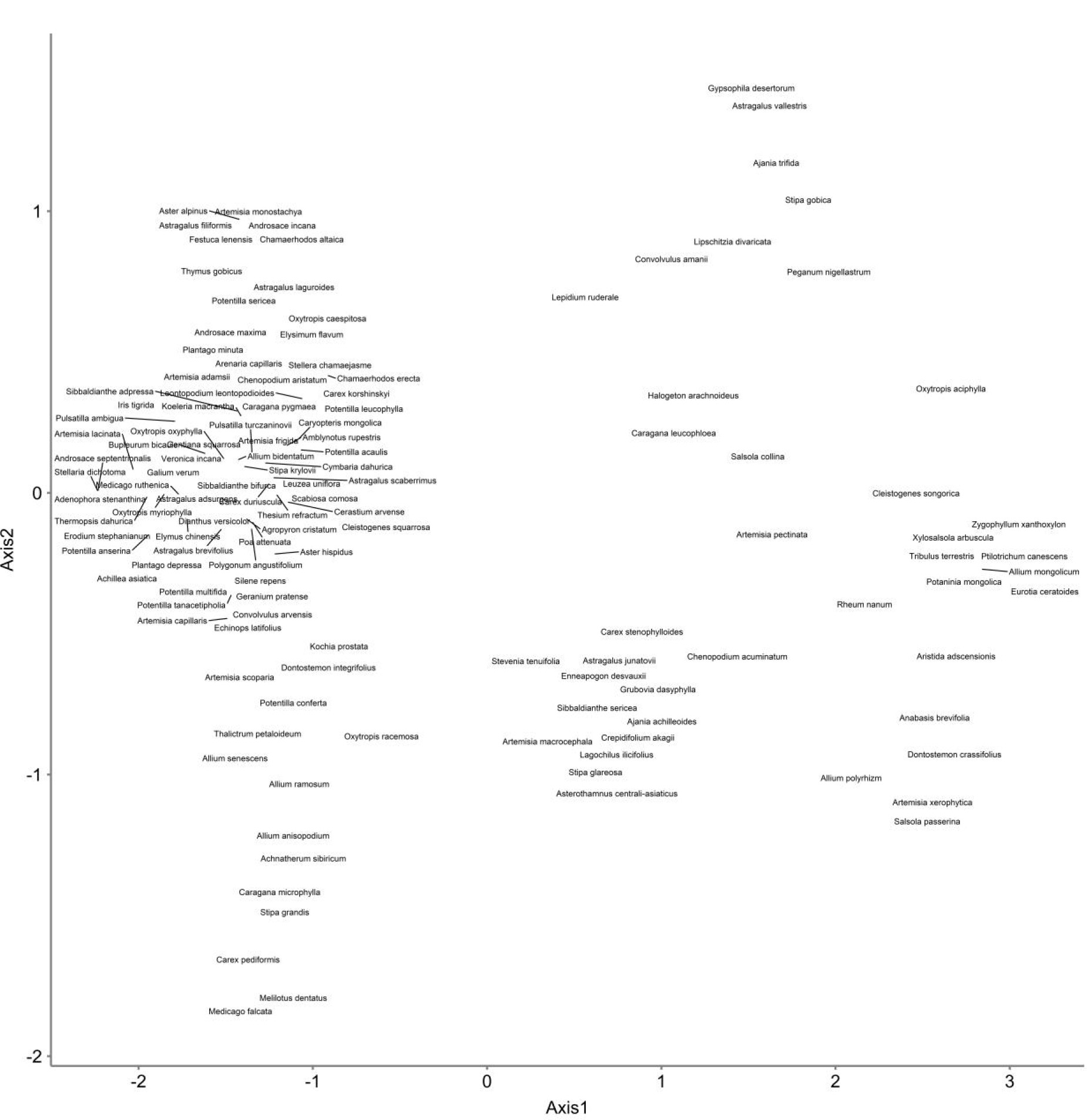
Result of loading values of species in nonmetric multi-dimensional scaling (NMDS) based on Bray-Curtis dissimilarity of ungrazed/grazed communities in the north and south regions.

**Figure S5.**
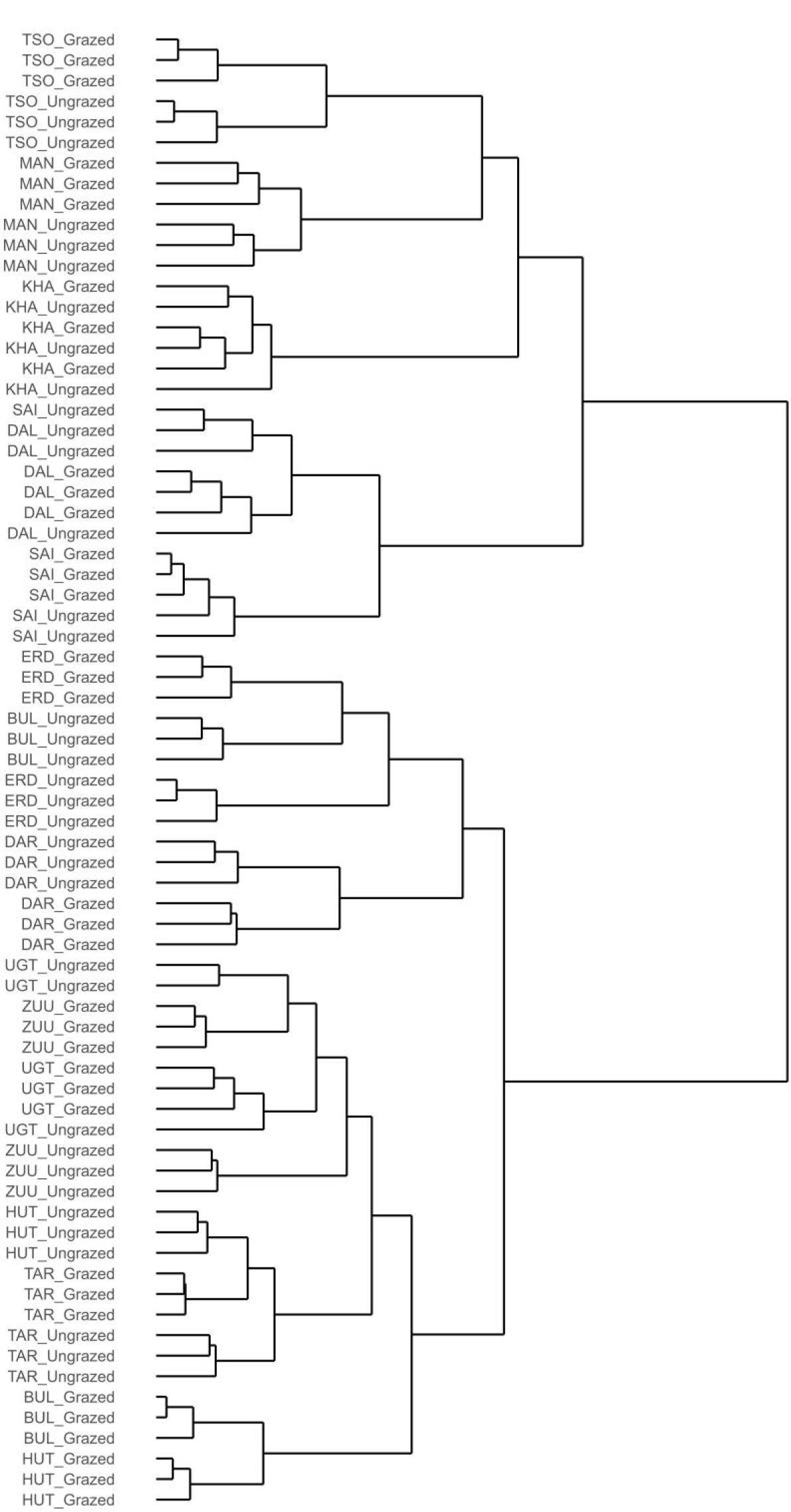
Result of clustering based on Ward’s algorithm of ungrazed/grazed communities in the north and south regions. Bray-Curtis dissimilarity matrix was used as a distance matrix.

**Table S1.** Partitioning of Bray-Curtis dissimilarity to balanced variation in abundance (species turnover) and abundance gradient (nestedness).

| Region | Ungrazed/grazed community | Balanced variation in abundance (%) | Abundance gradient (%) |
| --- | --- | --- | --- |
| North (semi-arid) region | Ungrazed community | 97.6 | 2.4 |
|  | Grazed community | 97.2 | 2.8 |
| South (arid) region | Ungrazed community | 95.4 | 4.6 |
|  | Grazed community | 91.4 | 8.6 |

**Table S2.** Mantel correlation coefficients based on Spearman’s ρ between variables in ungrazed and grazed communities of the north and south regions. Bold characters denote statistically significant correlations.

| Variable | Variable | North_Ungrazed | North_Grazed | South_Ungrazed | South_Grazed |
| --- | --- | --- | --- | --- | --- |
| Geographic distance | Aridity | **0.748** | **0.748** | **0.790** | **0.790** |
| Geographic distance | pH | 0.128 | **0.236** | 0.117 | **0.327** |
| Geographic distance | Sand | **0.188** | **0.275** | 0.137 | **0.460** |
| Geographic distance | Clay | 0.072 | -0.036 | -0.010 | -0.074 |
| Aridity | pH | 0.079 | **0.174** | 0.008 | **0.472** |
| Aridity | Sand | **0.336** | **0.237** | 0.171 | **0.255** |
| Aridity | Clay | -0.050 | 0.050 | 0.089 | -0.088 |
| pH | Sand | 0.037 | **0.176** | -0.009 | -0.030 |
| pH | Clay | **0.289** | -0.006 | -0.019 | 0.024 |
| Sand | Clay | **0.261** | 0.069 | **0.423** | 0.132 |

**Table S3.** Spearman’s ρ between aridity and relative abundance of shrub species in ungrazed and grazed communities of the north and south regions. Bold characters denote statistically significant correlations.

| Region | Fence | $\rho$ |
| --- | --- | --- |
| North (semi-arid)<br>region | Ungrazed<br>community | 0.295 |
|  | Grazed<br>community | 0.106 |
| South (arid)<br>region | Ungrazed<br>community | **0.622** |
|  | Grazed<br>community | **0.644** |

